# Multi-scale simulations of membrane adhesion mediated by CD47-SIRP*α* complexes

**DOI:** 10.1101/2025.01.22.633554

**Authors:** Ruihan Hou, Shuanglong Ren, Rong Wang, Bartosz Różycki, Jinglei Hu

## Abstract

Adhesion of biological cells is essential for various processes, including tissue formation, immune responses, and signaling. It involves multiple length scales, ranging from nanometers to micrometers, which are characteristic of (a) the intercellular receptor-ligand binding that mediates the cell adhesion, (b) the spatial distribution of the receptor and lignad proteins in the membranes of adhering cells, (c) adhesion-induced deformations and thermal undulations of the membranes, (d) the overall size of the interface between adhering cells. Therefore, computer simulations of cell membrane adhesion require multi-scale modeling and suitable approximations that capture the essential physics of the system under study. Here, we introduce such a multi-scale approach to study membrane adhesion mediated by the CD47-SIRP*α* binding, which is an immunologically relevant process. The synergetic use of coarse-grained molecular dynamics simulations and mesoscale kinetic Monte Carlo simulations allows us to explore both equilibrium properties and dynamical behavior of adhering membranes on the relevant length scales between 1 nm and 1 µm on time scales ranging from 0.1 ns all the way up to about 20 s. The multi-scale simulations not only reproduce available experimental data but also give quantitative predictions on binding-induced conformational changes of SIRP*α* and membrane-mediated cooperativity of the CD47-SIRP*α* binding as well as fluctuation-induced interactions between the CD47-SIRP*α* complexes. Our approach is applicable to various membrane proteins and provides invaluable data for comparison with experimental findings.

## 1 Introduction

Adhesion of biological cells arises from the specific binding of membrane receptors to their ligands anchored in the plasma membrane of an apposing cell, which is essential for various processes, including tissue formation, immune responses, and signaling. SIRP*α* receptors are present in the plasma membrane of macrophages [1, 2]. Their ligands are the ubiquitous ‘marker of self’ proteins CD47 (Fig. 1a). The specific binding of the SIRP*α* receptors to the CD47 ligands results in inhibition of engulfment of ‘self’ cells by macrophages and thus constitutes a key checkpoint of our innate immune system. Consequently, the CD47-SIRP*α* binding has been recognized as a potential therapeutic target in cancer and inflammation [2, 3].

**Figure 1:**
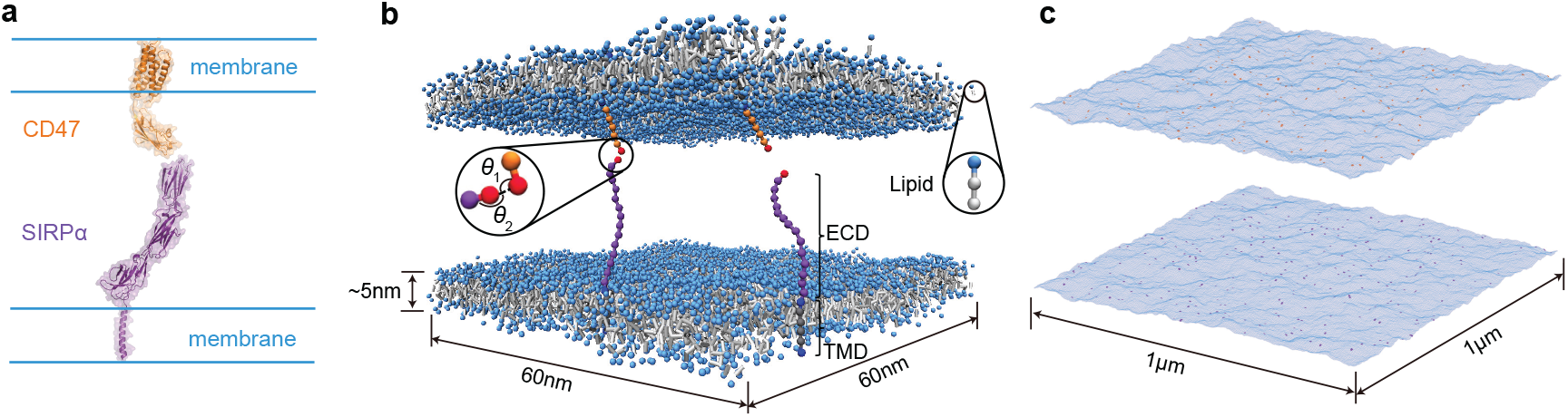
Illustration of the binding interaction between CD47 (orange) and SIRP*α* (purple) in the membrane (blue) environment. (a) Cartoon of the CD47-SIRP*α* complex constructed using available crystal structures of CD47 and SIRP*α*. In CD47 (PDB code 7MYZ), an extracellular domain (ECD) is connected to a transmembrane domain (TMD) via a short linker. SIRP*α* consists of three IgSF domains (PDB code 2WNG) and a TMD. The TMD is linked to one of the IgSF domains via a long polypeptide segment. (b) Snapshot from coarse-grained molecular dynamics simulations. The CD47 and SIRP*α* molecules are modeled by chains of beads and bind specifically *via* a potential that depends on the distance between the binding sites (in red) and the two angles *θ*_1_ and *θ*_2_ as depicted. Each of the lipid membranes has a thickness of about 5 nm and a projected area of 60 nm × 60 nm. (c) Snapshot from Monte Carlo simulations of a mesoscale lattice model. The membranes are modeled as discrete elastic surfaces that undergo thermal undulations. The CD47 and SIRP*α* molecules are modeled as single particles that diffuse along the membrane surfaces and bind to form the CD47-SIRP*α* complex when they are at opposite membrane sites and within an appropriate distance. The membranes here have the projected area of 1 µm^2^.

Physical processes involved in cell adhesion have been studied at different length and time scales using computer simulations. All-atom molecular dynamics simulations have been used to explore conformational fluctuations and concerted motions of single receptor molecules, such as the T cell receptor–CD3 complex embedded in a lipid membrane [4, 5]. Coarse-grained molecular simulations have been employed to study membrane-mediated cooperativity of receptor-ligand binding in systems with dozens of receptors and ligands anchored in lipid membranes with the lateral size of up to 100 nm [6]. Lattice-type models based on the Helfrich theory of membrane elasticity [7, 8, 9] have been used to simulate adhesion zones with the linear extension of up to several micrometers [10, 11, 12]. An appropriate combination of these methods should enable to study adhering membranes in a broad range of length and time scales.

The adhesion of cell membranes involves multiple length scales ranging from Angstroms to micrometers. Firstly, the specific receptor-ligand binding occurs on the length scale of Angstroms to nanometers. Secondly, the thickness of the cell membrane is about 5 nm. Thirdly, the extension of the extracellular domains of the receptors and ligands is typically of the order of 10 nm. Fourthly, the lateral distance between the receptor-ligand complexes involved, e.g. in immune responses or signaling is typically of the order of 100 nm. Finally, interfaces between adhered cells can be a few micrometers in size. Therefore, computer simulations of cell membrane adhesion require multi-scale modeling and suitable approximations that capture the essential physics of the system under study. Here, we introduce such a multi-scale approach to study membrane adhesion mediated by the CD47-SIRP*α* binding (Fig. 1), which is an immunologically relevant process. Firstly, we introduce an implicit-solvent coarse-grained molecular model for the system of membranes adhering by the CD47-SIRP*α* binding (Fig. 1b). We parametrize this coarse-grained model to reproduce available data from various experiments. Then we perform extensive molecular dynamics (MD) simulations to characterize both equilibrium and kinetic properties of the system under study. Next, we adapt a lattice-based mesoscale model and conduct kinetic Monte Carlo (MC) simulations of the CD47-SIRP*α* adhesion system (Fig. 1c). The MC simulations not only yield the CD47-SIRP*α* binding equilibrium and rate constants in quantitative agreement with the MD simulation results but also provide accurate predictions on indirect, fluctuation-induced, membrane-mediated interactions between the CD47-SIRP*α* complexes. The synergetic use of the coarse-grained MD simulations and the mesoscale MC simulations allows us to explore both equilibrium properties and dynamical behavior of adhering membranes on the relevant length scales between 1 nm and 1 µm (Fig. 1) and on time scales ranging from about 0.1 ns all the way up to about 20 s.

## 2 Results and Discussion

We adapted the implicit-solvent coarse-grained model of lipid bilayers introduced by Cooke and Deserno [13] to simulate a system of two membranes, where the lower membrane contained SIRP*α* receptors and the upper membrane comprised CD47 molecules (Fig. 1b). In the frame-work of this model, the extracellular domain (ECD) of SIRP*α* consist of 15 beads whereas the ECD of CD47 is formed of 6 beads. All the beads are taken to have the same diameter, *σ*_0_ = 1 nm, which sets the basic length scale of the coarse-grained model. The SIRP*α* receptors bind their ligands in the 1:1 stoichiometry and this binding gives rise to the adhesion of the two membranes. The CD47-SIRP*α* binding is caused by an attractive interaction between single beads at the tips of the ECDs of SIRP*α* and CD47 (these beads are marked in red in Fig. 1b). This attractive interaction depends on the distance between the two beads as well as on the local angle between the ECDs of SIRP*α* and CD47. The linear size of the ECDs as well as the geometry of the CD47-SIRP*α* binding are incorporated into the model based on the molecular structures available in the Protein Data Bank (PDB) under the accession codes 7MYZ and 2WNG [14, 15]. The trans-membrane domains (TMDs) of SIRP*α* and CD47 are formed of lipid-type-beads so that they are kept within the lipid bilayers in the course of the MD simulations. A detailed description of the coarse-grained model is given in Methods.

To determine the binding equilibrium constant of soluble variants of SIRP*α* and CD47, *K*_3D_, we performed MD simulations of only the ECDs of SIRP*α* and CD47 with no lipid membranes (Fig. 2a). We simulated several concentrations of the ECDs (i.e. several systems with different volumes and different numbers of pairs of the ECDs of SIRP*α* and CD47). For each of the concentrations, we conducted a relaxation run of 22.5 ms and a subsequent production run of up to 6.75 s, during which over 4000 binding and unbinding events were recorded. We assumed here that a CD47-SIRP*α* pair was in a bound state if the energy of interaction between their binding beads was below −2 *k*_B_*T*. We estimated *K*_3D_ using two alternative approaches. Firstly, we computed the binding equilibrium constant directly from the definition *K*_3D_=[CD47-SIRP*α*]/[SIRP*α*][CD47], where [SIRP*α*] and [CD47] denote the average concentration of free receptors and free ligands, respectively, whereas [CD47-SIRP*α*] is the concentration of receptor-ligand complexes. Secondly, we used the maximum likelihood analysis [6]. We found that both approaches gave consistent results with 1*/K*_3D_ = (1.8 ± 0.2) µM in the whole range of studied concentrations (Fig. 2a). This result is in very good agreement with the CD47-SIRP*α* dissociation constant values of 1 to 2 µM as measured in experiments and reported in the literature [2, 1].

**Figure 2:**
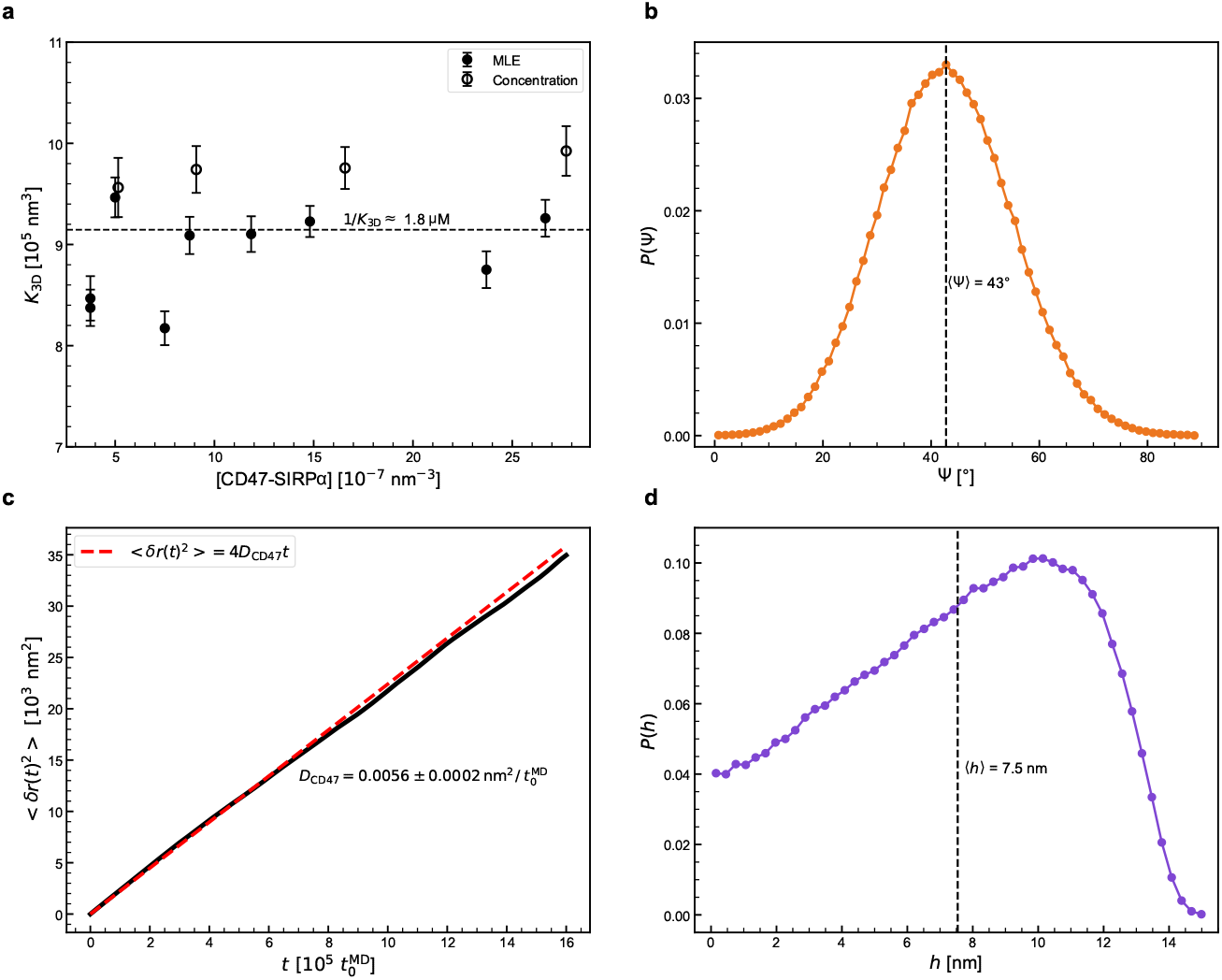
Validation of the coarse-grained simulation approach. (a) Binding equilibrium constant *K*_3D_ for the ECDs of CD47 and SIRP*α* in the bulk (i.e. in the absence of lipid membranes) versus concentration of the CD47-SIRP*α* complexes. The data points were obtained from simulations with different concentrations of CD47 and SIRP*α*. The filled circles indicate the *K*_3D_ values obtained from the maximum likelihood analysis [6] whereas the open circles correspond to the *K*_3D_ values computed directly from the definition *K*_3D_ = [CD47-SIRP*α*]*/*[CD47][SIRP*α*]. The average value of *K*_3D_ corresponds to the dissociation constant 1*/K*_3D_ ≈ 1.8 µM, consistent with the experimental range between 1 and 2 µM [1, 2]. (b) Distribution of angle Ψ between the CD47 ECD and the membrane plane. Results obtained from the simulations of a single membrane comprising three molecules of CD47. The dashed line indicates the average angle Ψ of 43^°^, which is consistent with 40^°^ obtained in all-atom molecular dynamics simulations [14]. (c) Mean squared displacement of the center-of-mass of the CD47 TMDs as a function of time. Results obtained from the simulations of a single membrane comprising three molecules of CD47. The red dashed line corresponds the least-square fit of the MD data to the Einstein relation ⟨*δr*(*t*)^2^⟩ = 4*D*_CD47_*t*, leading to the diffusion coefficient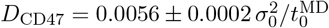. Matching this value to the CD47 diffusion coefficient of 0.125±0.02 µm^2^/s, as determined in fluorescence experiments with giant plasma membrane vesicles [10], determines the basic time unit in the MD simulations,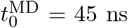. (d) Distribution of height *h* of the membrane-anchored SIRP*α*. Results obtained from the simulations of a single membrane comprising three molecules of SIRP*α*. The height *h* is defined as the shortest distance between the tip of the SIRP*α* ECD and the membrane plane. The dashed line indicates the average height of 7.5 nm, which is in agreement with the experimental value [16].

Then we performed MD simulations of one membrane comprising the full-length CD47 proteins and enclosed in a box of side lengths *L*_*x*_ = *L*_*y*_ = 30 *σ*_0_ and *L*_*z*_ = 50 *σ*_0_. The number of lipids in the membrane was adjusted in such a way that the membrane was under no mechanical tension. There were three CD47 molecules anchored in the membrane. The simulations had a length of 45 ms, during which the angle *ψ* between the membrane plane and the CD47 ECD was measured. The distribution of *ψ* was found to be approximately Gaussian with the average value of 43^°^ (Fig. 2b). This result is consistent with the equilibrium value of *ψ* = 40^°^ obtained by Fenalti et al. from all-atom MD simulations of the full-length CD47 [14] and shows that the coarse-grained MD simulations correctly capture the orientation of the ECD relative to the TMD.

In these MD simulations we also measured the two-dimensional diffusion coefficient *D*_CD47_ of the membrane-anchored CD47 molecules. Namely, we measured the mean squared displacement of the center-of-mass of the CD47 TMDs as a function of time, and fitted this dependence to the Einstein relation ⟨*δr*(*t*)^2^⟩ = 4*D*_CD47_*t* (Fig. 2c). By matching 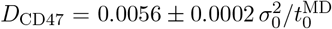 determined from the MD simulations to the value of 0.125±0.02 µm^2^*/*s measured in fluorescence experiments with giant plasma membrane vesicles [10], we obtained an estimate for the basic time unit in the MD simulations,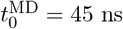.

Next we performed MD simulations of one membrane comprising the full-length SIRP*α* receptors. The simulation box was a cube of side lengths *L*_*x*_ = *L*_*y*_ = 30 *σ*_0_ and *L*_*z*_ = 50 *σ*_0_. The membrane contained three molecules of SIRP*α* and the number of lipids was adjusted in such a way that the membrane was under no mechanical tension. The simulation time was 45 ms. We measured the height *h* of SIRP*α*, i.e. the smallest distance between the tip of the SIRP*α* ECD and the membrane surface. We found that the distribution of *h* was rather broad with the average value of 7.5 nm (Fig. 2d). The same average height of SIRP*α* has been determined in cell surface optical profilometry experiments [16], which shows that the coarse-grained MD simulations properly capture the anchoring of SIRP*α* in the lipid bilayer.

The coarse-grained MD simulation results presented in Fig. 2 are consistent with available data from various experiments and all-atom MD simulations [2, 1, 16, 14]. Having validated the coarse-grained model for SIRP*α* and CD47, we performed MD simulations of a system of two membranes, where the lower membrane contained *N*_p_ receptors (SIRP*α*) and the upper membrane comprised *N*_p_ ligands (CD47). We carried out the MD simulations using a cubic box with dimensions *L*_*x*_ × *L*_*y*_ × *L*_*z*_ and periodic boundary conditions. We simulated four systems with (i) *L*_*x*_ = *L*_*y*_ = 15 *σ*_0_ and *N*_p_ = 2, (ii) *L*_*x*_ = *L*_*y*_ = 30 *σ*_0_ and *N*_p_ = 4, (iii) *L*_*x*_ = *L*_*y*_ = 60 *σ*_0_ and *N*_p_ = 5, and (iv) *L*_*x*_ = *L*_*y*_ = 90 *σ*_0_ and *N*_p_ = 8. In each of the four systems, the height of the simulation box was *L*_*z*_ = 100 *σ*_0_ and the number of lipids was adjusted in such a way that the membranes were under no mechanical tension. The MD trajectories had a length of up to 1.8 ms.

In the MD simulations of the largest system, i.e. system (iv), we measured how the ECDs of CD47 (Fig. 3a) and SIRP*α* (Fig. 3d) were oriented relative to the membrane. More precisely, we measured the angle between the ECD and the membrane normal, both in the bound (filled dots in Fig. 3) and unbound (open dots in Fig. 3) states of CD47 and SIRP*α*. We found that the distribution of orientations of the CD47 ECD was unaffected by the CD47-SIRP*α* binding (Fig. 3a). However, the distribution of the SIRP*α* ECD orientations was much broader in the unbound state than in the bound state (Fig. 3d), demonstrating that the binding to CD47 imposes restraints on the orientation of SIRP*α* relative to the membrane.

**Figure 3:**
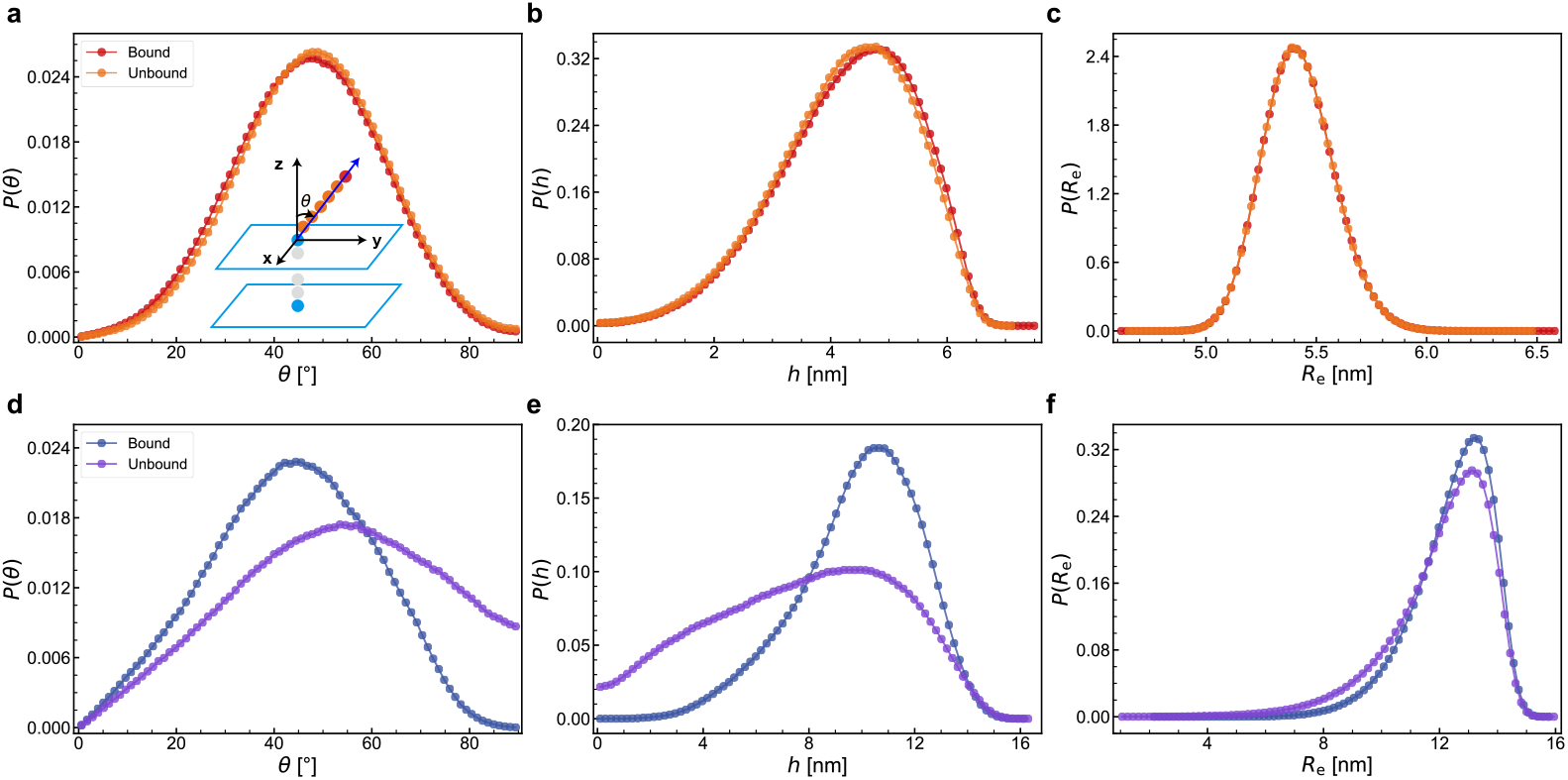
Conformational statistics of CD47 (a-c) and SIRP*α* (d-f) in the bound (filled circles) and unbound (open circles) states as obtained from the coarse-grained MD simulations of the two-membrane system. (a,d) Distribution of the orientation angle *θ* measured between the membrane normal and the ECD of CD47 (a) or SIRP*α* (d). (b,e) Distribution of the molecular height *h* measured from the membrane plane to the ECD tip of CD47 (b) or SIRP*α* (e). (c,f) Distribution of the end-to-end distance *R*_e_ of the ECD of CD47 (c) or SIRP*α* (f).

In the MD simulations of system (iv) we also measured the height *h* of the CD47 (Fig. 3b) and SIRP*α* (Fig. 3e) molecules, both in the bound and unbound states. We defined the height *h* as the smallest distance between the ECD tip and the membrane surface. We found that the distribution of CD47 height was unaffected by the CD47-SIRP*α* binding (Fig. 3b). However, the distribution of SIRP*α* height was clearly narrower in the bound state than in the unbound state (Fig. 3e), indicating that the binding to CD47 makes the ECD of SIRP*α* effectively stiffer.

In the MD simulations of system (iv) we also measured the end-to-end distance of the ECDs of CD47 (Fig. 3c) and SIRP*α* (Fig. 3f), both in the bound (filled dots in Fig. 3) and unbound (open dots in Fig. 3) states. We found the distribution of the CD47 ECD end-to-end distance to be insensitive to the CD47-SIRP*α* binding (Fig. 3c). However, the distribution of the SIRP*α* ECD end-to-end distance was noticeably narrower in the bound state than in the unbound state, showing that the conformations of the SIRP*α* ECD are affected by the CD47-SIRP*α* binding.

Taken together, the simulation results shown in Fig. 3 demonstrate that SIRP*α* changes its orientations and conformations upon binding to CD47. It is tempting to suggest that such changes can be a means of transferring a ‘do-not-eat-me’ signal from ‘self’ cells to macrophages.

For each of the four systems, (i)-(iv), we determined the thermal roughness *ξ*_⊥_ of the adhering membranes, which was computed as the variance of the equilibrium distribution of the local distance between the membranes. We also determined the two-dimensional binding constant *K*_2D_ using the maximum likelihood analysis of binding and unbinding events observed in each of the four MD trajectories [6]. We found the dependence of *K*_2D_ on *ξ*_⊥_ to follow the generic relationship derived by Hu et al. [6]

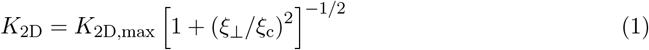

with *ξ*_c_ representing a confinement length. The best fit of the simulation data to Eq. (1) was obtained for parameters *K*_2D,max_ = 95000 ± 900 nm^2^ and *ξ*_c_ = 2.50 ± 0.06 nm (Fig. 4b).

**Figure 4:**
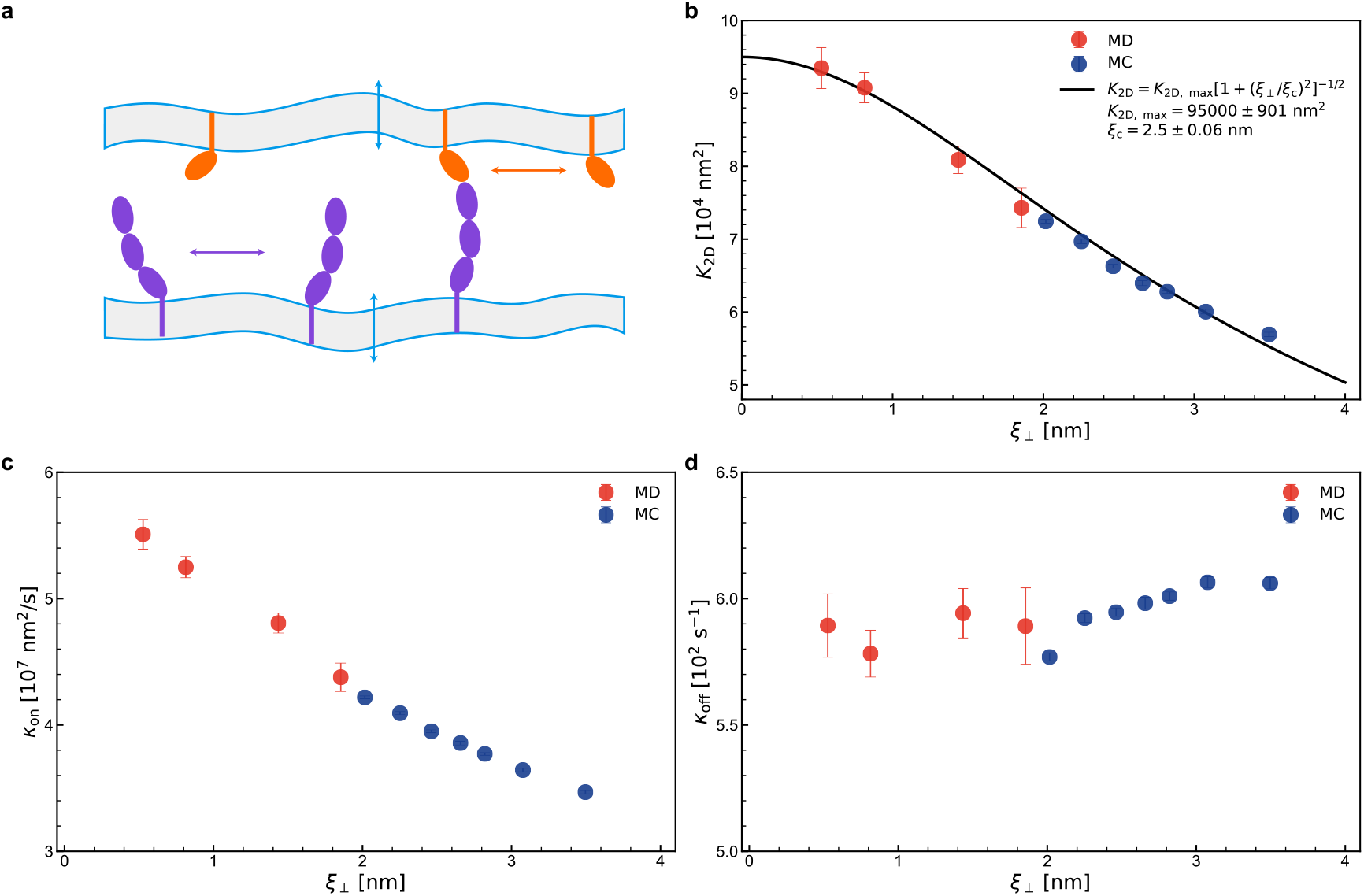
Membrane-mediated binding cooperativity. (a) Schematic of trial moves in the MC simulations. The blue arrows indicate thermal undulations of the membranes. The orange and purple arrows represent the lateral diffusion of CD47 and SIRP*α* molecules along the membrane surface. (b) Binding equilibrium constant *K*_2D_, (c) on-rate constant *k*_on_, and (d) off-rate constant *k*_off_ *versus* the relative membrane roughness *ξ*_⊥_ obtained from the coarse-grained MD (points in red) and MC simulations (points in blue). The solid line in (b) represents the least-square fit to all the data points.

The relationship between *K*_2D_ and *ξ*_⊥_ as given by Eq. (1) reflects a positive cooperativity in the CD47-SIRP*α* binding process, which can be explained as follows: Fluid membranes are rather soft and undergo thermal fluctuations. The formation CD47-SIRP*α* complexes suppresses membrane fluctuations and causes the average distance between the membranes to be closer to the optimal distance for the CD47-SIRP*α* binding, which in turn facilitates the formation of additional CD47-SIRP*α* complexes between the two membranes. The feedback between the suppression of membrane fluctuations and the formation of receptor-ligand complexes leads to an effect of membrane-mediated binding cooperativity, which has been predicted theoretically [17], examined in dissipative particle dynamics (DPD) simulations of a generic coarse-grained model [6], and confirmed quantitatively in fluorescence microscopy experiments with GFP-tagged CD47 on giant plasma membrane vesicles binding to SIRP*α* immobilized on a surface [10].

The maximum likelihood analysis allowed us to determine also the on- and off-rate constants, *k*_on_ and *k*_off_, for the CD47-SIRP*α* binding. While *k*_on_ was found to decrease with *ξ*_⊥_ (Fig. 4c), *k*_off_ was practically constant, independent of *ξ*_⊥_ (Fig. 4d). The latter result is in contrast with the weak dependence of *k*_off_ on *ξ*_⊥_ reported by Hu et al. [6], probably because the relatively fast off-rates in the DPD simulations were not reaction-limited.

As mentioned before, we performed MD simulations of four systems of different sizes. The largest of the four systems had the lateral size of 90 nm and comprised eight molecules of CD47 in the upper membrane and eight molecules of SIRP*α* in the lower membrane. To study even larger systems, we adapted a mesoscale lattice-type model and employed kinetic MC simulations.

The mesoscale model is based on the Helfrich theory of membrane elasticity and represents membranes as discrete surfaces with the lattice size *a* = 5 nm comparable to the membrane thickness (Fig. 1c). Any site in the lower membrane can accommodate only one receptor (SIRP*α*) and any site in the upper membrane can be occupied by only one ligand (CD47). The receptors and ligands are represented in this model as single particles with no internal degrees of freedom (Fig. 1c). To ensure the specific CD47-SIRP*α* binding, one receptor only binds one ligand if two conditions are fulfilled: firstly, both the receptor and the ligand are located at opposite membrane sites, and secondly, the local distance *l* between these two opposite sites is within a binding range, i.e., 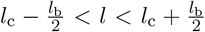, where *l*_c_ is the extension of the receptor-ligand complex and *l*_b_ is the width of the binding potential. In the MC simulations we used *l*_c_ = 3 *a* = 15 nm and *l*_b_ = 1.4 *a* = 7 nm. The CD47-SIRP*α* binding energy was *U*_b_ = 8 *k*_B_*T*. The bending rigidity of each of the two membranes was taken to be 13 *k*_B_*T*, corresponding to coarse-grained lipid membranes in the Cooke-Deserno model [13]. A detailed description of the mesoscale lattice-type model and of the kinetic MC simulation algorithm is given in Methods.

We performed the MC simulations with different numbers *N*_p_ = 20, …, 500 of CD47-SIRP*α* pairs and different lattice sizes, ranging from 20 × 20 to 200 × 200. We determined both the thermal roughness *ξ*_⊥_ and the two-dimensional binding constant *K*_2D_ using the same methods as in the analysis of the MD trajectories. Importantly, the dependence of *K*_2D_ on *ξ*_⊥_ obtained from the MC simulations was found to follow the same relationship given by Eq. (1) with *K*_2D,max_ = 95000 ± 900 nm^2^ and *ξ*_c_ = 2.50 ± 0.06 nm as the dependence of *K*_2D_ on *ξ*_⊥_ determined from the coarse-grained MD simulations (Fig. 4b). This result indicates that the mesoscale lattice-type model is properly parameterized to quantitatively reproduce the membrane-mediated binding cooperativity observed in the coarse-grained MD simulations.

MC simulations with local trial moves can be used to study membrane dynamics in the overdamped limit [18, 19]. From the MC trajectories we determined also the binding rate constants, *k*_on_ and *k*_off_, using the maximum likelihood method. Taking the MC time unit 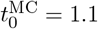 µs, the on- and off-rate constants from the MC simulations were found to be in good quantitative agreement with those obtained from the coarse-grained MD simulations (Figs. 4c and 4d). This result means that the MC simulations with local trial moves properly capture not only the equilibrium but also the kinetics of the CD47-SIRP*α* binding (Fig. 4). It is also worth noting that the MC time unit 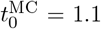 µs is more than an order of magnitude larger than the MD time unit, 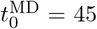 ns, and a MC simulation run comprising 2 × 10^7^ cycles, as performed in this study, corresponds to the physical time of about 22 s.

In order to quantify the indirect, fluctuation-induced, membrane-mediated interactions between the CD47-SIRP*α* complexes, we performed MC simulations of four systems that differed in size (200 × 200 or 400 × 400 lattice sites) and in the number of CD47-SIRP*α* pairs (*N*_p_ = 200 or 400). The overall area concentrations of CD47 and SIRP*α* in the four systems were *c*_0_ = 50, 100, 200, and 400 µm^−2^ (Fig. 5). For each of the systems, five independent parallel simulations were performed to measure the radial distribution function *g*(*r*) (Fig. 5b), the potential of mean force *U* (*r*) = −*k*_B_*T* ln(*g*(*r*)) (Fig. 5c), and the two-dimensional second virial coefficient 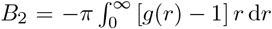 (Fig. 5d). At any of the protein concentrations studied here, *g*(*r*) *>* 1 and *∂g/∂r <* 0 (Figs. 5b), or equivalently *U* (*r*) *<* 0 and *∂U/∂r >* 0 (Figs. 5c), indicating an effective attraction between the CD47-SIRP*α* complexes. The effective attraction is rather weak (|*U* (*r*)| *< k*_B_*T* in the range of protein concentrations studied here) and has a long range. In addition, both the magnitude and the range of the effective attraction decrease with increasing *c*_0_ (Figs. 5b and 5c), which can be understood as follows: As the protein concentration is increased, the average distance between the CD47-SIRP*α* complexes decreases while the thermal undulations of the membranes get suppressed, thereby limiting the range and magnitude of the effective, membrane-mediated attraction.

**Figure 5:**
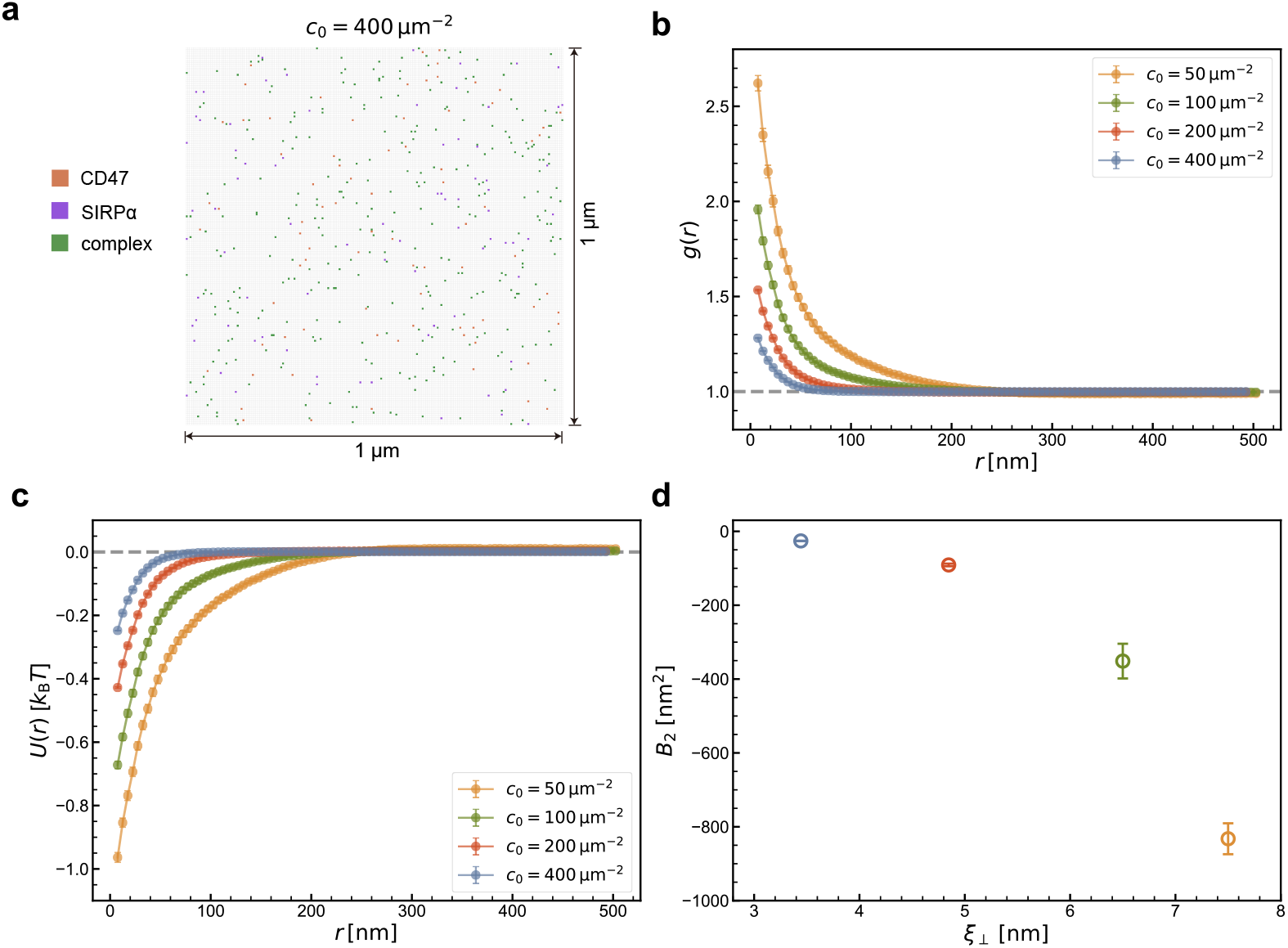
Membrane-mediated interactions between the CD47-SIRP*α* complexes as obtained from the MC simulations. (a) Simulation snapshot (top view) of a lattice with 200 × 200 sites, corresponding to a membrane segment with the surface area of 1 µm^2^. The free CD47 and SIRP*α* proteins are marked as dotes in orange and purple, respectively. The CD47-SIRP*α* complexes are marked as green dots. The protein concentration is *c*_0_ = 400 µm^−2^. (b) Pair correlation function *g*(*r*) of the CD47-SIRP*α* complexes at different protein concentrations, *c*_0_ = 50, 100, 200, and 400 µm^−2^. (c) Potential of mean force *U* (*r*) = −*k*_B_*T* ln(*g*(*r*)) at the different protein concentrations. (d) Two-dimensional second virial coefficient 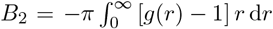 as a function of the membrane roughness *ξ*_⊥_.

The negative values of the second virial coefficient *B*_2_ (Fig. 5d) not only confirm that the effective interactions between the CD47-SIRP*α* complexes are attractive but also quantify the prepotency of the CD47-SIRP*α* complexes to cluster. As the membrane roughness *ξ*_⊥_ increases, *B*_2_ becomes more negative, implying that the effective attraction of the CD47-SIRP*α* complexes is indeed induced by membrane fluctuations (Fig. 5d). Taken together, even though the the effective membrane-mediated interactions between CD47-SIRP*α* complexes are rather weak (|*U* (*r*)| *< k*_B_*T*), the large-scale MC simulations provide accurate predictions on the range and magnitude of these interactions, which is practically impossible to achieve in the coarse-grained MD simulations.

## 3 Models and Methods

### Coarse-grained molecular model

We have adapted the Cooke-Deserno model of lipid bilayers [13] to simulate the membrane proteins CD47 and SIRP*α*. Each CD47 or SIRP*α* molecule consists of a transmembrane domain (TMD) and an extracellular domain (ECD) that includes extracellular beads (PE, in orange or purple) and a binding site (PB, in red) (Fig. 1b). Specifically, one CD47 consists of 12 beads and one SIRP*α* of 21 beads. The TMD of a CD47 or SIRP*α* is composed of 6 beads, featuring four hydrophobic lipid-tail-like beads (PT, in dark gray) between two lipid-head-like beads (PH, in dark blue). Each lipid molecule consists of one ‘head’ bead (LH) and two ‘tail’ beads (LT). The hard-core repulsion between any pair of two beads are modeled by the purely repulsive potential

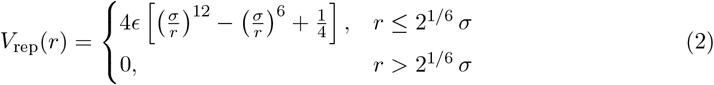

where *σ* = *σ*_0_ and *ϵ* = *ϵ*_0_ for most of the pairs. Here, *σ*_0_ is the basic length unit and *ϵ*_0_ the basic energy unit. For LH-LH and LH-LT pairs, *σ* = 0.95 *σ*_0_. For PT-LH and PH-LT pairs, *ϵ* = 10 *ϵ*_0_. To ensure the CD47-SIRP*α* binding with 1:1 stoichiometry, *σ* = 3.5 *σ*_0_ and *ϵ* = 10 *ϵ*_0_ are chosen for PB-PB pairs that belong to two CD47 or SIRP*α* molecules.

Adjacent beads within the CD47 or SIRP*α* molecules are connected via the harmonic potential

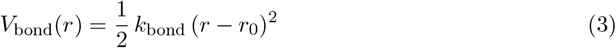

with the spring constant 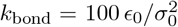 and the rest length *r*_0_ = *σ*_0_. In each lipid, the LH and the last LT beads are also bonded via this potential with 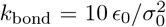 and *r*_0_ = 4 *σ*_0_. Additionally, any two consecutive beads of each lipid are linked by the finite extensible nonlinear elastic (FENE) bond

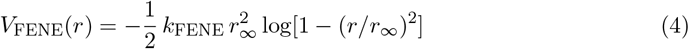

with the stiffness 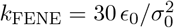 and the divergence length *r*_∞_ = 1.5 *σ*_0_.

The lipid tail beads experience pairwise attractive potentials

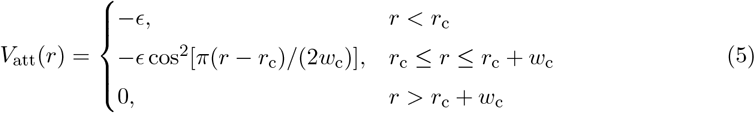

where *r*_c_ = 2^1*/*6^ *σ*_0_, *w*_c_ = 1.6 *σ*_0_, and *ϵ* = *ϵ*_0_ are chosen to obtain a stable fluid bilayer with the bending modulus of about 13 *k*_B_*T* at room temperature [13], which is within the bending rigidity range of about 10 to 40 *k*_B_*T* determined from experimental measurements of lipid bilayers in a fluid state [20, 21]. This attraction effectively accounts for the hydrophobic interactions of the lipid molecules in the Cooke-Deserno implicit solvent model. Any pair of a lipid tail bead and a lipid-tail-like bead in CD47 or SIRP*α*, i.e. PT-LT pairs, also interact through this potential to facilitate the insertion of the TMDs of CD47 and SIRP*α* into the lipid bilayer.

To capture the conformational flexibility of CD47 and SIRP*α* molecules as well as the orientation of their ECDs relative to the membrane, every three adjacent beads in each CD47 or SIRP*α* interact via the bending potential

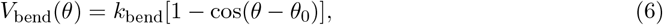

where *k*_bend_ is the strength and *θ*_0_ the preferred angle. For the TMD beads of CD47 or SIRP*α, k*_bend_ = 100 *ϵ*_0_ and *θ*_0_ = 180^°^ are set to maintain a rigid linear structure of TMDs within the lipid bilayers [22]. Since CD47 consists of one single ECD [14, 23] and a SIRP*α* consists of three ECDs connected by linkers [24, 25], *k*_bend_ = 100 *ϵ*_0_ and *k*_bend_ = 10 *ϵ*_0_ are chosen for their ECD beads, respectively. The ECDs of CD47 and SIRP*α* are stable and modeled as rigid units, whereas the linkers between adjacent ECDs can be flexible. For the ECD beads of CD47 or SIRP*α, θ*_0_ = 180^°^. For PE-PH-PT of each CD47, *k*_bend_ = 100 *ϵ*_0_ and *θ*_0_ = 130^°^ are set according to the results of all-atom molecular dynamics simulations which show that the CD47 ECD forms an angle of approximately 50^°^ with the membrane normal [14]. For PE-PH-PT of each SIRP*α, k*_bend_ = 10 *ϵ*_0_ and *θ*_0_ = 140^°^, leading to an average height of 7.5 nm measured between the membrane plane and the ECD tip, in good agreement with experimental results [16].

The specific binding of CD47 and SIRP*α* is modeled via the distance- and angle-dependent potential

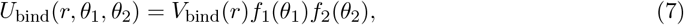

where the radial part *V*_bind_(*r*) is given by

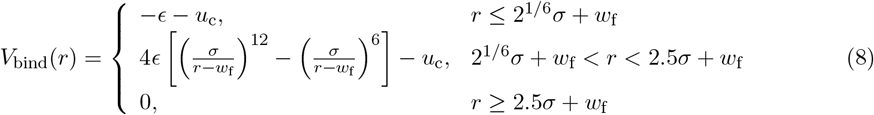

with *σ* = *σ*_0_, *w*_f_ = 0.4 *σ*_0_, *ϵ* = 22 *ϵ*_0_ and *u*_c_ = 4 *ϵ*_0_ [(1*/*2.5)^12^ −(1*/*2.5)^6^] ≈ −0.016 *ϵ*_0_. The angular part *f*_*i*_(*θ*_*i*_) with *i* = 1, 2 takes the form

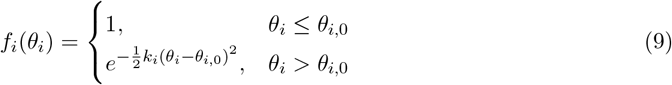

with *θ*_1,0_ = 90^°^, *θ*_2,0_ = 0^°^ and *k*_1_ = *k*_2_ = 10 rad^−2^. The angles *θ*_1_ and *θ*_2_ are defined by the two binding sites and their adjacent PE beads, as illustrated in Fig. 1b. Our choice of the parameters yields a dissociation constant of about 1.8 µM for soluble CD47 and SIRP*α* molecules that lack their TMD, consistent with the experimental range of 1 to 2 µM [1, 2].

Taken together, the force field parameters of the coarse-grained MD model are adjusted to effectively account for the length and flexibility of the protein ECDs, orientation of the proteins relative to the membrane, and the specific receptor-ligand binding. Therefore, our coarse-graining protocol is transferrable to other membrane proteins and their complexes.

### 3.2 Molecular dynamics simulations

MD simulations of the coarse-grained model introduced in the previous subsection were performed using the Python GPU-Accelerated Molecular Dynamics software (PYGAMD) [26]. MD runs in the canonical ensemble were conducted within a cuboid box of side lengths *L*_*x*_, *L*_*y*_ and *L*_*z*_ under periodic boundary conditions. A constant temperature *T* = 1.1 *ϵ*_0_*/k*_B_ was maintained using a Langevin thermostat with a drag coefficient 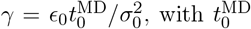 being the basic time unit and *k*_B_ denoting the Boltzmann constant.

Four membrane systems with *L*_*x*_ = *L*_*y*_ = 15 *σ*_0_, *L*_*x*_ = *L*_*y*_ = 30 *σ*_0_, *L*_*x*_ = *L*_*y*_ = 60 *σ*_0_ and *L*_*x*_ = *L*_*y*_ = 90 *σ*_0_ were simulated. The corresponding numbers of pairs of CD47 and SIRP*α* were *N*_p_ = 2, 4, 5, and 8. The extension of the simulation box in the direction perpendicular to the membranes was *L*_*z*_ = 100 *σ*_0_ in all of the four systems. Initially, the two membranes were pre-assembled in such a way that they were both planar and parallel to the *x*-*y* plane of the rectangular simulation box. The distance between the lower surface of the upper membrane and the upper surface of the lower membrane (in other words, the distance between the mid-planes of the two membranes minus the membrane thickness) was chosen to be 23 nm, i.e. slightly larger than the sum of lengths of the fully-extended ECDs of CD47 and SIRP*α*, which is 21 nm. In equilibrium, the average separation between the two adhering membranes was found to be between 14.3 and 14.8 nm (Fig. S1), i.e. clearly smaller than the initial separation of 23 nm. The number of lipids in each of the membranes was adjusted in such a way that the membranes were under no mechanical tension. Matching the lipid bilayer thickness of about 5 *σ*_0_ to the experimental value of 5 nm led to *σ*_0_ ≈ 1 nm. Comparing the diffusion coefficient *D* = 0.125 µm^2^*/*s of CD47 in the cell membrane [10] to our simulation result 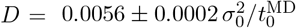 (Fig. 2c) led to the time unit 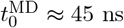. The diffusion coefficient of the lipids was found to be 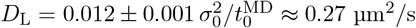 (Fig. S2). The integration time step was set to 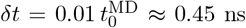. For each of the four systems, a relaxation run of 2 × 10^7^ *δt* ≈ 9 ms was performed for thermal equilibration and a subsequent run of up to 4 ×10^9^ *δt* ≈ 1.8 s was conducted for statistical sampling. From these simulations, approximately 4000 binding and unbinding events are observed in each of the simulated systems. The binding rate constants and the equilibrium constants were extracted from the MD trajectories using the maximum likelihood method as described below.

We also performed MD simulations of a single membrane containing three CD47 molecules enclosed within a cubic box of size *L*_*x*_ = *L*_*y*_ = 30 *σ*_0_ and *L*_*z*_ = 50 *σ*_0_. 25 independent runs, each of 45 ms, were conducted to measure the diffusion coefficient of CD47. We extracted the center-of-mass coordinates of the CD47 TMDs from the trajectories and calculated the mean squared displacement ⟨*δr*(*t*)^2^⟩ as a function of time *t* to obtain the data shown in Fig. 2c. The least-square fit of the data to the Einstein relation ⟨*δr*(*t*)^2^⟩ = 4*Dt* yielded the diffusion coefficient 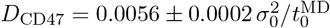.

To measure the CD47-SIRP*α* binding constant in the bulk, i.e. in the absence of lipid membranes, four systems with ECDs of CD47 and SIRP*α* were simulated in a cubic box of side length *L*. The simulated molecules lacked their TMDs. In these systems, 10 or 15 pairs of ECDs of CD47 and SIRP*α* were enclosed in a cubic box of side length *L* = 150 *σ*_0_ or *L* = 200 *σ*_0_. Each of the four systems was subject to a relaxation run of 5 × 10^7^ *δt* ≈ 22.5 ms and a production run of up to 15 × 10^9^ *δt* ≈ 6.75 s for data acquisition, during which over 4000 binding and unbinding events were recorded to determine the binding constant.

### 3.3 Mesoscale lattice-type model for membrane adhesion

In the coarse-grained MD simulations described in the previous subsection, the maximum membrane size was 90×90 nm^2^. Extending the simulations to larger spatial and temporal scales necessitates significant computational resources. To efficiently handle these larger scales, we adopt a lattice model, which offers a more computationally feasible approach [19, 9, 11, 27]. In this model, the membranes are described as discrete surfaces (blue and grey in Fig. 1c). To capture the whole spectrum of bending deformations of the flexible membranes, the size of each quadratic patch on the lattice is chosen to be *a* = 5 nm to match the membrane thickness [28]. For two tensionless membranes that have no spontaneous curvature and are on average parallel, the bending energy is given by the discretized Helfrich Hamiltonian

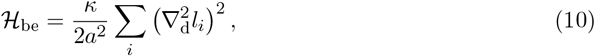

where 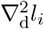 is the discrete Laplacian of the local separation *l*_*i*_ of the two membranes at lattice site *i*. The two membranes cannot penetrate each other, implying *l*_*i*_ *>* 0 for all the lattice sites. The separation field {*l*_*i*_} specifies the relative configuration of the two adhering membranes.

In Eq. (10), *κ* = *κ*_1_*κ*_2_*/*(*κ*_1_ + *κ*_2_) is the effective bending rigidity of the two membranes with bending moduli *κ*_1_ and *κ*_2_. The bending rigidity of the lipid membranes used in the coarse-grained MD simulations is *κ*_1_ = *κ*_2_ = 13 *k*_B_*T* [13]. Thus the effective bending rigidity *κ* = 6.5 *k*_B_*T* is chosen in the lattice model to match the coarse-grained MD model.

The CD47 and SIRP*α* molecules are modeled as particles which occupy single vacant membrane patches (in orange and purple), diffuse along the membrane surfaces and bind specifically to form inter-membrane complexes with 1:1 stoichiometry, as illustrated in Fig. 1c. More than one CD47 or SIRP*α* molecule in each membrane are not allowed to occupy the same membrane patch in order to account for intermolecular hardcore repulsion. The distribution of CD47 on the upper membrane is described by the compositional variable 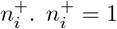 indicates that the upper membrane patch at lattice site *i* is occupied by a CD47, whereas 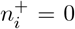 indicates the membrane patch contains no CD47. Likewise, the distribution of SIRP*α* on the lower membrane is described by bivalent variables 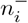 with 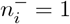 and 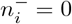 indicating the presence and absence of a SIRP*α* on the lower membrane patch at lattice site *i*, respectively. The adhesion energy due to CD47-SIRP*α* binding takes the form

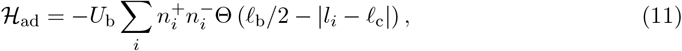

where the binding is characterized by a square-well potential centered at *ℓ*_c_ with strength *U*_b_ and width *ℓ*_b_. *ℓ*_c_ is the average length of the CD47-SIRP*α* complex. The Heaviside step function Θ(…) requires that the local membrane separation *l*_*i*_ should be within the binding range *ℓ*_c_ −*ℓ*_b_*/*2 *< l*_*i*_ *< ℓ*_c_ +*ℓ*_b_*/*2 for the complex to form. Eq. (11) implies that a CD47 binds to a SIRP*α* only if the two molecules occupy the apposing membrane patches at the same lattice site and the local separation of the two membrane patches is within the binding range. Despite its simplicity, the square-well potential effectively takes into account the distance and orientation dependence of CD47-SIRP*α* binding.

The three parameters *U*_b_, *ℓ*_b_ and *ℓ*_c_ of the binding potential in Eq. (11) are determined according to the coarse-grained MD simulation results. The maximal binding constant in the lattice model here is 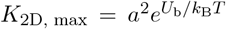 when the two membranes are flat and rigid and their separation is within the binding range. Matching the coarse-grained MD result *K*_2D, max_ = 95000 ± 900 nm^2^ leads to *U*_b_ ≈ 8 *k*_B_*T*. The width of the binding potential *ℓ*_b_ = 1.4 *a* = 7 nm is chosen to reproduce the dependence of the binding constant *K*_2D_ on the relative thermal roughness *ξ*_⊥_ of the two membranes, *K*_2D_ = *K*_2D, max_[1 + (*ξ*_⊥_*/ξ*_c_)^2^]^−1*/*2^, as measured in the coarse-grained MD simulations and firstly derived by Hu et al [6]. The roughness *ξ*_⊥_ is defined as 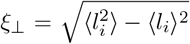 with ⟨… ⟩ denoting the ensemble average. The length scale *ξ*_c_ = 2.50 ± 0.06 nm reflects intrinsic variations in the extension of the CD47-SIRP*α* complex in the direction perpendicular to the membranes. The smaller *ξ*_c_, the stronger the restriction of the local membrane separation imposed by the complexes. The average length of the complexes *ℓ*_c_ = 15 nm is determined by the average separation of the two adhering membranes in the coarse-grained MD simulations (Figs. S1). In this model, the lengths of CD47 and SIRP*α* are *ℓ*_CD47_ = 5 nm and *ℓ*_SIRP*α*_ = 10 nm according to their average heights in the complexes from the membranes as obtained from coarse-grained MD simulations (Figs. 3c and 3g). These molecular lengths determine the minimal separation of two apposing membrane patches when either or both of the patches accommodate the molecules.

### 3.4 Monte Carlo simulations

MC simulations of the lattice model were carried out using our in-house C++ code[29, 11]. There were two types of trial moves implemented in the MC simulations: (i) horizontal translations of the particles representing the membrane proteins, and (ii) local vertical displacements of each of the discrete surfaces representing the membranes. In a MC move of type (i), one CD47 or SIRP*α* was randomly selected and shifted from its original position at site *i* to one of the four nearest neighbor sites *j*. If site *j* was already occupied by another CD47 or SIRP*α*, or if the local membrane separation *l*_*j*_ at site *j* was smaller than the height of the selected protein, the trial move was rejected. If the selected CD47 or SIRP*α* formed an inter-membrane complex at site *i*, an attempt was made to either break the complex with probability 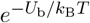 or move the complex as a whole with probability 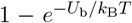 provided that site *j* on the opposite membrane was vacant and *ℓ*_c_ − *ℓ*_b_*/*2 *< l*_*j*_ *< ℓ*_c_ + *ℓ*_b_*/*2. In a MC move of type (ii), the local separation of the two membranes at a randomly selected site *i* was changed to 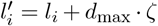, with *d*_max_ = 0.2 *a* = 1 nm being the maximum displacement and *ζ* representing a random number distributed uniformly between −1 and 1. When the membranes at site *i* were both occupied, this trial move might trigger the formation or breaking of a CD47-SIRP*α* complex, depending on the values of *l*_*i*_ and 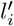. Any trial move of type (ii) that led to penetration of the two membranes or to insertion of CD47 or SIRP*α* into the opposite membrane was rejected. The probability of accepting any trial move was determined from the detailed balance condition by using the Metropolis algorithm with Hamiltonian given by Eqs. (10) and (11).

In one MC cycle, all membrane sites were on average attempted to be displaced vertically once and all the CD47 and SIRP*α* molecules were on average attempted to be shifted horizontally once. We found that the MC simulations produced the on- and off-rate constants consistent with those obtained from the coarse-grained MD simulations.

The MC simulations were performed with a square lattice of *N* × *N* sites with *N* = 20, 30, 40, 60, 100, 150 and 200. The corresponding numbers of protein molecules in each of the membranes were *N*_p_ = 20, 30, 40, 60, 200, 300 and 400. For each of the systems, a relaxation run of 10^5^ MC cycles was performed for thermal equilibration, followed by up to 2 × 10^7^ MC cycles for statistical sampling. The on- and off-rate constants were determined using the maximum likelihood method (described in the following subsection) from the sequences of binding and unbinding events identified in MC simulations.

To quantify membrane-mediated interactions between the CD47-SIRP*α* complexes, we performed additional MC simulations of four systems with *N* = 200 or *N* = 400 and *N*_p_ = 200 or *N*_p_ = 400. The overall area concentrations of CD47 and SIRP*α* in these four systems were *c*_0_ = 50, 100, 200, and 400 µm^−2^. Five independent parallel simulations were performed for each of the systems. Each of these simulations comprised 2 × 10^6^ MC cycles for thermal equilibration and up to 10^7^ MC cycles for statistical sampling. From these series of simulations, the pair correlation function *g*(*r*) of the CD47-SIRP*α* complexes was determined at each of the protein concentrations.

### 3.5 Maximum likelihood estimation of *k*_on_, *k*_off_, and *K*_2D_

Here, we briefly review the maximum likelihood estimation developed by one of the authors for extracting the binding kinetics from simulation trajectories [6], and apply this method to both the MD and MC simulations. In the coarse-grained MD simulations, a CD47-SIRP*α* pair is assumed to be in the bound state if the energy of interaction between their binding sites is lower than −2 *k*_B_*T*. Otherwise, the two molecules are taken to be in the unbound state. In the MC simulations, a CD47-SIRP*α* pair is considered to be in the bound state if the two molecules are located at opposite membrane sites and if their distance is within the range of *ℓ*_c_ −*ℓ*_b_*/*2 *< l*_*j*_ *< ℓ*_c_ +*ℓ*_b_*/*2. A pair of CD47 and SIRP*α* is in the unbound state if they do not have the appropriate location or separation. With these definitions, the binding and unbinding events are identified based on the bound and unbound states of the CD47 and SIRP*α* molecules in a given simulation trajectory. These binding and unbinding events divide each of the simulation trajectories into time windows, each of which is characterized by the number of CD47-SIRP*α* complexes.

A system with *N*_CD47_ ligands and *N*_SIRP*α*_ receptors has totally (*N* + 1) states, where *N* = min(*N*_CD47_, *N*_SIRP*α*_) is the maximum number of CD47-SIRP*α* complexes. The simulation trajectories can be mapped to a Markov model

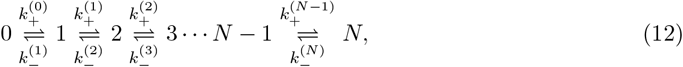

where the transition rates 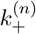 and 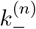 are, respectively, related to the on- and off-rate constants 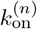 and 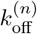 via

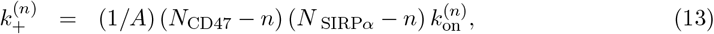

and

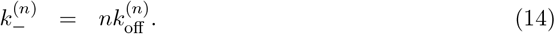

The equilibrium constant that characterizes the CD47-SIRP*α* binding affinity is defined as

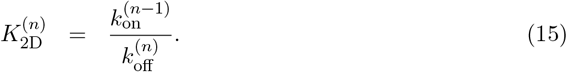

The on- and off-rate constants 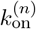 and 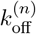 in Eqs. (13)-(14) can be determined from the observed numbers of transitions between the states and from the overall dwell times in the states. The binding and unbinding events divide the simulation trajectories into time windows *i* of length *t* in state *n*_i_, which are followed by a transition into state *n* + *s* with *s*_i_ = 1 or −1. The probability for staying in state *n*_*i*_ for a dwell time 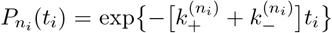. The probability for the time window *i* with the observed transition is 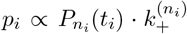 for *s*_*i*_ = 1 and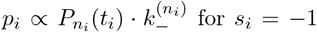. The likelihood function is the probability of the whole trajectory and takes the form

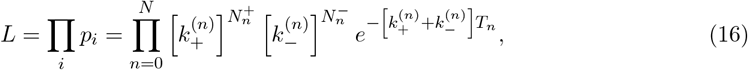

where 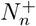 is the total number of transitions from state *n* to *n* + 1, 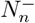 the total number of transitions from state *n* to *n* − 1, and *T*_*n*_ the total dwell time in state *n*.

Maximizing the likelihood function *L* in Eq. (16) with respect to the rate constants 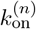 and 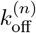 leads to the maximum likelihood estimators

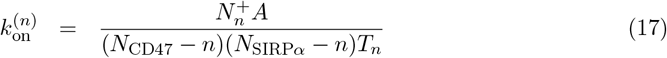

And

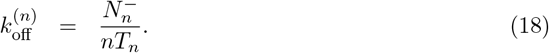

Our estimator for the binding constant defined in Eq. (15) is

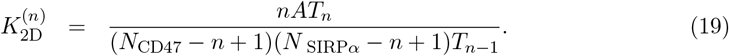

In each trajectory, generated in either the coarse-grained MD simulations or the mesoscale MC simulations, we identify the transition numbers 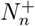 and 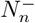 as well as the overall dwell times *T*_*n*_ in each state, and estimate the constants 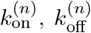 and 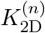 according to Eqs. (17)-(19). Next, for each of the simulated systems, we collect equilibrium configurations with the same number *n* of CD47-SIRP*α* complexes and compute the thermal roughness 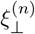 of the membranes based on these configurations. Then the equilibrium constant 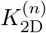 and rate constants 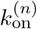 and 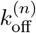 are associated with 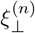 for each *n* = 0, …, *N*. Each data point in Fig. 4 represents one simulated system with that value of *n* which corresponds to the largest number of binding and unbinding events.

## 4 Conclusions

In our multi-scale modeling framework, we incorporated coarse-grained MD and mesoscale MC methods to simulate membrane adhesion mediated by the CD47-SIRP*α* binding. Thanks to the use of GPU-accelerated software [26], we were able to carry out extensive MD simulations of the adhering membranes with the lateral size of up to 90 nm, reaching the time scale of 1 s. Using the mesoscale lattice-based model, we performed kinetic MC simulations of even larger systems with the lateral size of up to 1 µm on time scales of up to 20 s. Importantly, we constructed the coarse-grained molecular model to accurately capture the geometry and flexibility of the CD47 and SIRP*α* proteins. We tuned parameters of the coarse-grained model to reproduce data from several independent experiments (Fig. 2). In addition, by carefully adjusting MC parameters, we matched the equilibrium and rate constants of the CD47-SIRP*α* binding to those obtained from the coarse-grained MD simulations (Fig. 4). These efforts together allowed us to explore the binding-induced conformational changes of SIRP*α* (Fig. 3), the membrane-mediated cooperativity of the CD47-SIRP*α* binding (Fig. 4), and the indirect, membrane-mediated, long-range attraction between the CD47-SIRP*α* complexes (Fig. 5). Our approach is applicable to various membrane proteins and facilitates access to detailed information on membrane adhesion at length scales ranging from 1 nm to 1 µm and time scales up to 10 s, providing invaluable data for comparison with experimental findings. Our multi-scale approach can also be applied to cell adhesion mediated by several types of receptor-ligand complexes, as present e.g. in the immunological synapse, and provide a physical picture for how different types of protein complexes form spatio-temporal patterns within the adhesion zone.

In the fluorescence microscopy experiments conducted by Steinkühler et al., GFP-labeled CD47 molecules on giant plasma membrane vesicles were observed to bind GST-tagged SIRP*α* molecules immobilized on a planar surface [10]. The two-dimensional binding constant *K*_2D_ was found to increase monotonically with the area concentration of the CD47-SIRP*α* complexes, indicating a positive cooperativity of the binding process. The binding cooperativity was attributed to positive feedback between formation of receptor-ligand complexes and suppression of thermal undulations of the adhering membranes, as predicted earlier on the basis of membrane elasticity theory and statistical mechanics [17, 6]. Here, we simulated the binding of membrane-anchored SIRP*α* and CD47 molecules. We found *K*_2D_ to decrease monotonically with membrane roughness *ξ*_⊥_ according to Eq. (1), in agreement with the effect of membrane-mediated binding cooperativity predicted by Hu et al. [6]. More experimental research is required to directly test the validity of Eq. (1) at physiological conditions. In particular, experiments with membrane-anchored SIRP*α* and CD47 molecules are needed to examine whether Eq. (1) with *K*_2D,max_ = 95000 ± 900 nm^2^ and *ξ*_c_ = 2.50 ± 0.06 nm, as found in the multi-scale simulations reported here (Fig. 4), correctly captures the CD47-SIRP*α* cooperative binding.

Another outcome of the coarse-grained MD simulations that could be examined in future experiments pertains the binding-induced conformational changes of SIRP*α*: When SIRP*α* binds CD47, it becomes effectively stiffer and attains a particular range of orientations with respect to the adhered membranes (Fig. 3). We suggest that such binding-induced changes in SIRP*α* conformations may be a means of transferring a ‘do-not-eat-me’ signal from ‘self’ cells to macrophages. A detailed picture of SIRP*α* conformations before and after the binding to CD47 could, in principle, be obtained from all-atom MD simulations. However, even with the use of state-of-the-art GPU-accelerated software, it would be extremely time consuming or even unfeasible to simulate on relevant time scales and in all-atom details such a large molecular system as the CD47-SIRP*α* complex in the environment of adhered membranes.

The MC simulations quantified the fluctuation-induced, membrane-mediated, long-ranged attraction between the CD47-SIRP*α* complexes (Fig. 5), providing predictions of the collective behavior of the CD47-SIRP*α* complexes in a native-like membrane environment. We argue that this kind of weak, membrane-mediated attraction between membrane proteins cannot be obtained from experiments alone.

Our results together demonstrate a critical role of membrane fluctuations and molecular flexibility in governing the binding cooperativity and dynamics of the CD47-SIRP*α* complexes, providing insights into mechanisms underlying immune checkpoint functions. Typically, MD simulations for the design of immunotherapeutic drugs focus primarily on the binding site [30]. While this approach provides important structural information about direct interactions between the drug and its target protein, it overlooks the conformational variability of the entire protein and the influence of membrane fluidity and fluctuations on the binding affinity and drug efficacy. Such simplifications may cause faulty predictions of the drug effectiveness, especially in complex biological environments. Future work could focus on more detailed protein conformational dynamics and on cis interactions between CD47 and SIRP*α* which possibly modulate their trans interactions [31].

## 5 Acknowledgements

This research was financially supported by the National Natural Science Foundation of China (grant number 22161132012, 22173045) and by the National Science Centre of Poland (grant number 2021/40/Q/NZ1/00017). The numerical simulations were performed using the computing facilities in the High-Performance Computing Center (HPCC) of Nanjing University and in the Centre of Informatics – Tricity Academic Supercomputer and Network (CI TASK) in Gdansk, Poland. For the purpose of open access, the authors have applied a CC-BY public copyright license to any Author Accepted Manuscript (AAM) version arising from this submission.

## NOMENCLATURE

*σ*_0_: length unit in the coarse-grained model
*ϵ*_0_: energy unit in the coarse-grained model
*V*_bond_: harmonic potential
*k*_bond_: spring constant
*V*_FENE_: finite extensible nonlinear elastic potential
*k*_FENE_: stiffness of FENE
*r*_∞_: divergence length
*V*_att_: pairwise attractive potential
*V*_bend_: bending potential
*k*_bend_: bending strength
*θ*_0_: preferred angle
*U*_bind_: binding potential
*V*_bind_: distance-dependent term of the binding potential
*f*_*i*_: angle-dependent term of the binding potential
*L*_*x*_, *L*_*y*_: lateral dimensions of simulation box
*L*_*z*_: height of simulation box
*k*_B_: Boltzmann constant
*γ*: drag coefficient
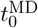: basic time unit in MD
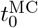: basic time unit in MC
*A*: membrane projected area
*N*_p_: numbers of CD47 or SIRP*α*
*δt*: integration time step
ℋ_be_: bending energy
*κ*: effective bending rigidity
*A*: grid spacing
*l*_*i*_: the local separation of the two membranes at lattice site *i*
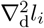: discrete Laplacian of *l*_*i*_
{*l*_*i*_}: separation field
ℋ_ad_: adhesion energy
*U*_b_: receptor-ligand binding energy
*ℓ*_b_: width of the binding potential
*ℓ*_c_: average length of the CD47-SIRP*α* complex
Θ(…): Heaviside step function
*n*_*i*_: occupation number showing CD47 presence/absence at lattice site *i* in upper membrane
*m*_*i*_: occupation number showing SIRP*α* presence/absence at lattice site *i* in lower membrane
*K*_2D_: two-dimensional binding constant
*K*_3D_: three-dimensional binding constant
*K*_2D, max_: maximal binding constant
*k*_on_: on-rate constant
*k*_off_: off-rate constant
*ξ*_⊥_: relative thermal roughness
*ξ*_c_: SD for fluctuations of *l*_*i*_ within the harmonic constraint
*d*_max_: maximum vertical displacement of the lattice
*ζ*: random number uniformly distributed between -1 and 1
*ψ*: angle between the CD47 ECD and the membrane plane
*h*: height between the binding site of CD47 and the membrane plane
*θ*: orientation angle of the ECD of CD47 or SIRP*α* relative to the membrane normal
LH: ‘head’ bead of lipid molecule
LT: ‘tail’ bead of lipid molecule
PE: extracellular bead of CD47 or SIRP*α*
PB: binding site of CD47 or SIRP*α*
PT: hydrophobic lipid-tail-like beads
*N*: maximum number of CD47-SIRP*α* complexes
*n*: number of CD47-SIRP*α* bonds
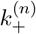: rate for the transition from state with *n* bonds to state with *n* + 1 bonds
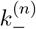: rate for the transition from state with *n* bonds to state with *n* − 1 bonds
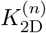: two-dimensional equilibrium constant for the binding reaction *n* − 1 ⇌ *n*
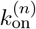: on-rate constant for the formation of the (*n*+1)-th bond at state *n*
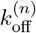: off-rate constant for the breaking of the *n*-th bond at state *n*
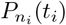: probability of staying in state *n*_*i*_ for a dwell time *t*_*i*_
*p*_*i*_: probability for the time window *i* with the observed transition
*L*: likelihood function
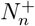: total number of transitions from state *n* to *n* + 1
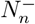: total number of transitions from state *n* to *n* − 1
*T*_*n*_: total dwell time in state *n*

