## Supplementary material for "Multi-scale simulations of membrane adhesion mediated by CD47-SIRP*α* complexes": Fig. S1, Fig. S2

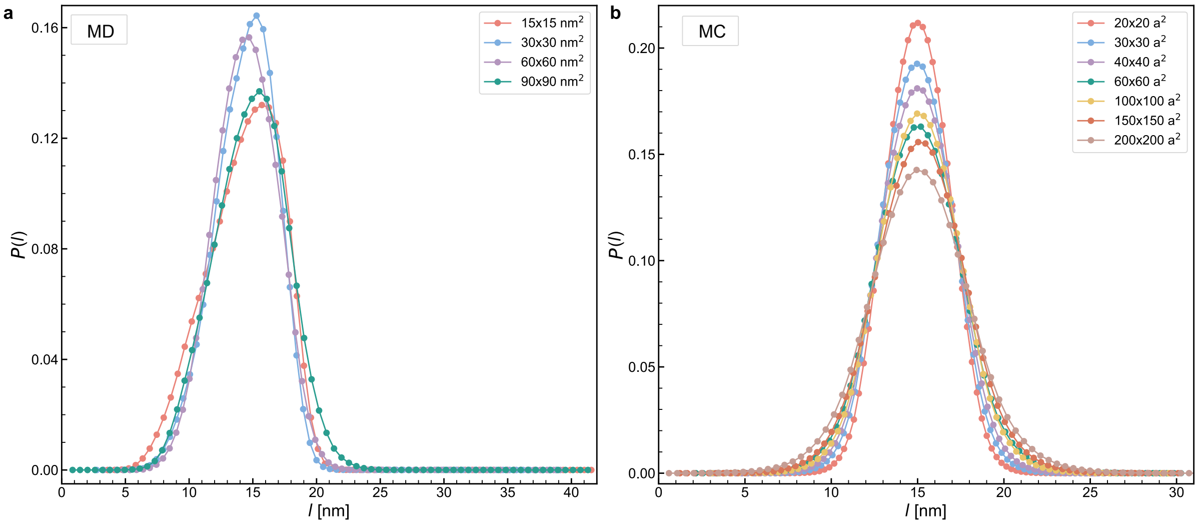


Figure S1: Distributions of the local membrane separation, $P(l)$, obtained from the coarse-grained MD (a) and MC (b) simulations with different membrane areas as specified in the legend.

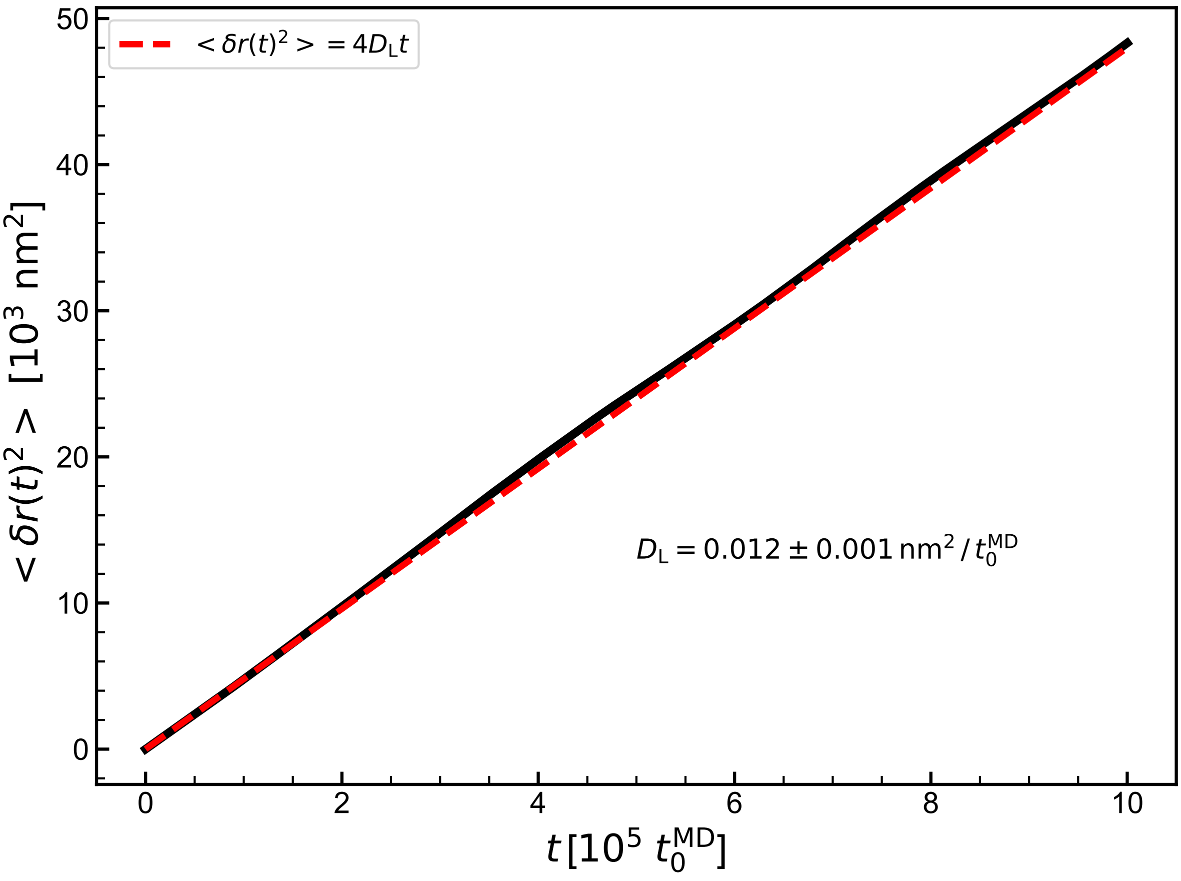


Figure S2: Mean squared displacement of the lipids as a function of time. The red dashed line corresponds the least-square fit of the MD data to the Einstein relation $\langle \delta r(t)^2 \rangle = 4D_\mathrm{L}t$, leading to the diffusion coefficient $D_\mathrm{L} = 0.012 \pm 0.001\,\mathrm{nm}^2/t_0^{\mathrm{MD}}$.
